# GenoSiS: A Biobank-Scale Genotype Similarity Search Architecture for Creating Dynamic Patient-Match Cohorts

**DOI:** 10.1101/2024.11.02.621671

**Authors:** Kristen Schneider, Murad Chowdhury, Mariano Tepper, Jawad Khan, Colorado Center for Personalized Medicine, Jonathan A. Shortt, Chris Gignoux, Ryan Layer

## Abstract

Many patients do not experience optimal benefits from medical advances because clinical research does not adequately represent them. While the diversity of biomedical research cohorts is improving, ensuring that individual patients are adequately represented remains challenging. We propose a new approach, GenoSiS, which leverages machine learning-based similarity search to dynamically find patient-matched cohorts across different populations quickly. These cohorts could serve as reference cohorts to improve a range of clinical analyses, including disease risk score calculations and dosage decisions. While GenoSiS focuses on finding genetic similarity within a biobank, our similarity search architecture can be extended to represent other medically relevant patient characteristics and search other biobanks.

---

Healthcare providers make treatment decisions by blending their experience, patient concerns, and data from clinical trials. While clinical trials are the gold standard for determining the safety and efficacy of medical interventions, health disparities arise, in part, because it is challenging to apply the results from a study to patients who are not represented by the subjects within the study cohort. Ding et al.^1^ quantified the genetic component of this effect by showing that the more distant an individual is from samples in the reference cohort, the less accurate their predicted disease risk was. Their results also showed that the straightforward approach of replicating a study for different groups (e.g., by genetic ancestry) provides only a marginal improvement for those who fit neatly into a group and no benefit for those who do not.

Mitigating health inequalities resulting from insufficient representation in clinical trial cohorts may necessitate creating reference cohorts tailored to individual patients. These *patient-matched cohorts* could, for example, be used to replicate clinical studies to generate evidence that inform representative treatment recommendations for all patients, especially those from populations that have traditionally been marginalized in healthcare research. While university biobanks are approaching the size and diversity required to fulfill many patients’ needs, traditional indexing and exact search methods do not scale. We propose leveraging machine learning-based approximate similarity search architectures to find cohorts of similar individuals dynamically.

Similarity search architectures find the nearest neighbors of an object using numerical vectors called embeddings. Each object is represented with an embedding, and objects with similar attributes have embeddings that are close in the vector space. A common approach to map objects to embeddings is to use a neural network trained with a corpus of objects with known distances. While embeddings are highly general and have been used for objects from words and video to DNA sequences, the primary challenge is determining the most relevant features. For diagnosis and treatment decisions, patient similarity is complex and context-dependent. Here, we start with genetic similarity, because it is stable, and genetic distance is a well-researched topic, but in the future, patient similarity will likely include other more dynamic data types, including methylation, expression, and health records, all of which can be represented as embeddings and can be explored using a similarity search database. Alternatively, structured fields can be incorporated to the search using hybrid techniques that include attribute filtering.

Our Genotype Similarity Search (GenoSiS) uses a novel genotype embedding model and the Intel^®^ Scalable Vector Search (SVS, https://github.com/intel/ScalableVectorSearch) performance library^2^ to find representative cohorts quickly for any patient. First, we produce positional encodings for each sample’s haplotype from the phased genotypes in a Variant Call File (VCF)^3^ and a recombination map^4^ (**Figure 1A**). Next, using a Siamese neural network ^5^, we learn genetic embeddings from pairs of haplotype encodings (**Figure 1B**), then produce embeddings for all haplotypes with the trained model and index them using SVS (**Figure 1C**). Using a sample’s genotypes as a query, GenoSiS can find the nearest neighbors with SVS, from which it constructs a sample’s matched cohort (**Figure 1D**). While existing methods^6–10^ offer static clusters and NxN similarity metrics, using SVS eliminates the need to recompute significant portions of the matrix when adding new samples. SVS also supports additional embeddings and future GenoSiS versions will represent non-genetic factors crucial for building cohorts for different diseases (e.g., methylation, medical records, etc.). We also plan to incorporate additional biobanks through a federated learning approach to address the risk of insufficient patient diversity within a single institution’s biobank. This method expands the pool of potential patient matches while minimizing patient-level data sharing, as it allows institutions to train models locally and share only the aggregated model parameters, reducing risks to patient privacy. GenoSiS software is available for download at https://github.com/kristen-schneider/precision-medicine.

**Figure 1.**
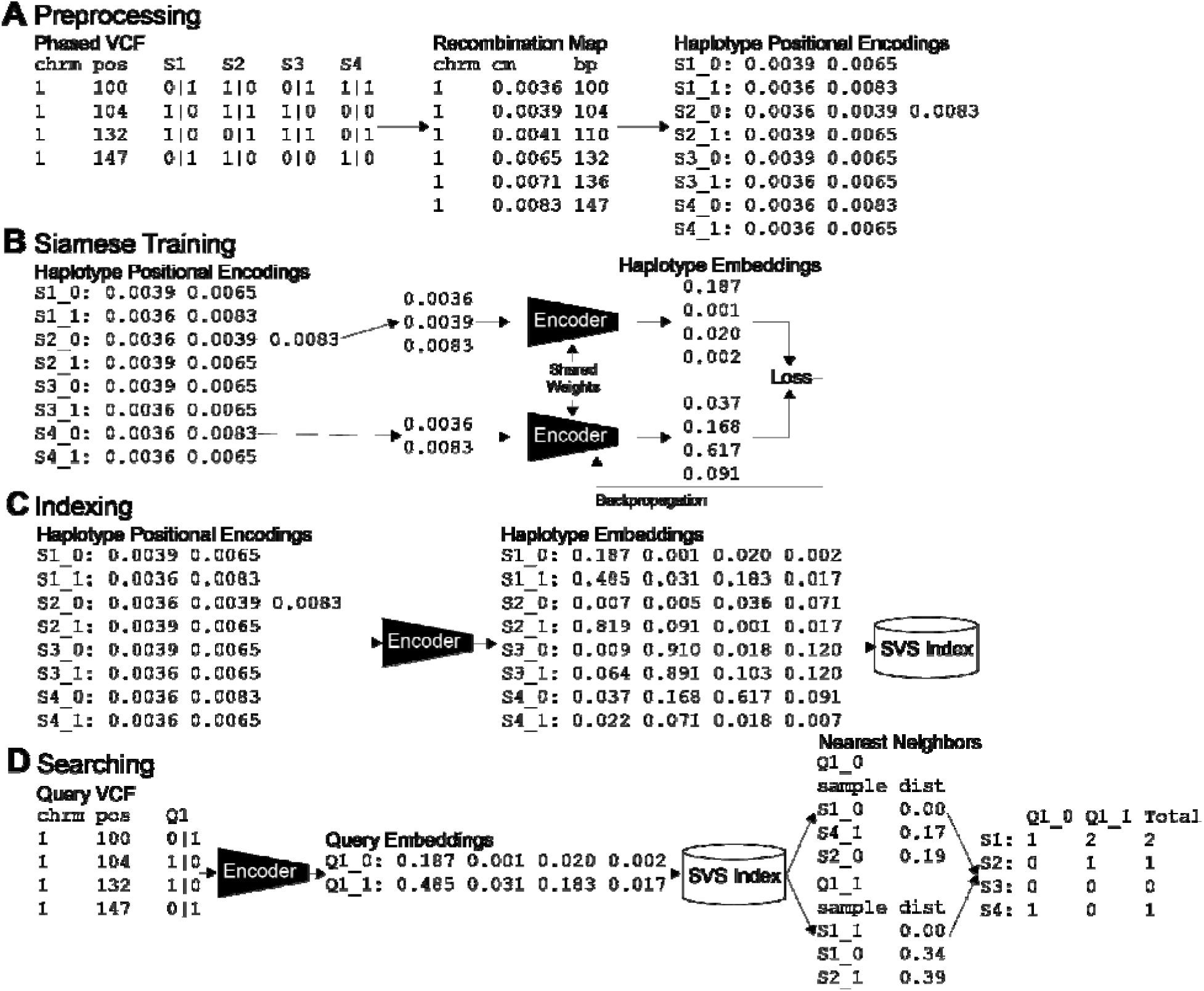
The Genotype Similarity Search (GenoSiS) workflow. **A** Starting with a VCF and recombination map, GenoSiS generates a haplotype positional encoding vector for each sample. **B** We then train an embedding model to minimize the difference between encoding and embedding distances. **C** Using the trained model, we create embeddings for all sample haplotypes and index them with Intel’s SVS. **D** The index is queried using a sample’s embeddings to find its nearest neighbors, and those results are aggregated to determine the sample’s cohort.

## II. RESULTS

GenoSiS quickly and accurately found high-quality matched cohorts across datasets and populations. In less than a second, GenoSiS identified representative cohorts among families in deCODE pedigrees, across populations among the 3,201 samples from the Thousand Genomes Project (TGP)^11–14^, and in the 73,346 samples from the Colorado Center for Personalized Medicine’s (CCPM) biobank^15^. GenoSiS is also highly general. With an embedding model trained on genotypes from whole genome sequencing, GenoSiS accurately constructed matched cohorts in CCPM data, which used targeted sequencing and imputation. These results demonstrate that GenoSiS is a robust method that has the potential to significantly improve patient representation in healthcare.

### A. Cohort Genetic Similarity

Genetic similarity can seem straightforward. We are most related to our parents and children, followed by siblings, grandparents, and so on. Outside our family, we are most related to those with similar geographic ancestry. Quantifying genetic similarity is a complex and active area of research^16^, and when we compare nominal and quantitative genetic similarity, the distributions can be surprisingly broad, overlapping, and potentially contradictory (**Figure 2**). With this uncertainty, there is no single genetic similarity ground truth from which we can measure accuracy and precision. To determine how well GenoSiS’s cohorts relate to the query sample, we compared our result to other widely used methods at different scales of relatedness and in different populations.

**Figure 2.**
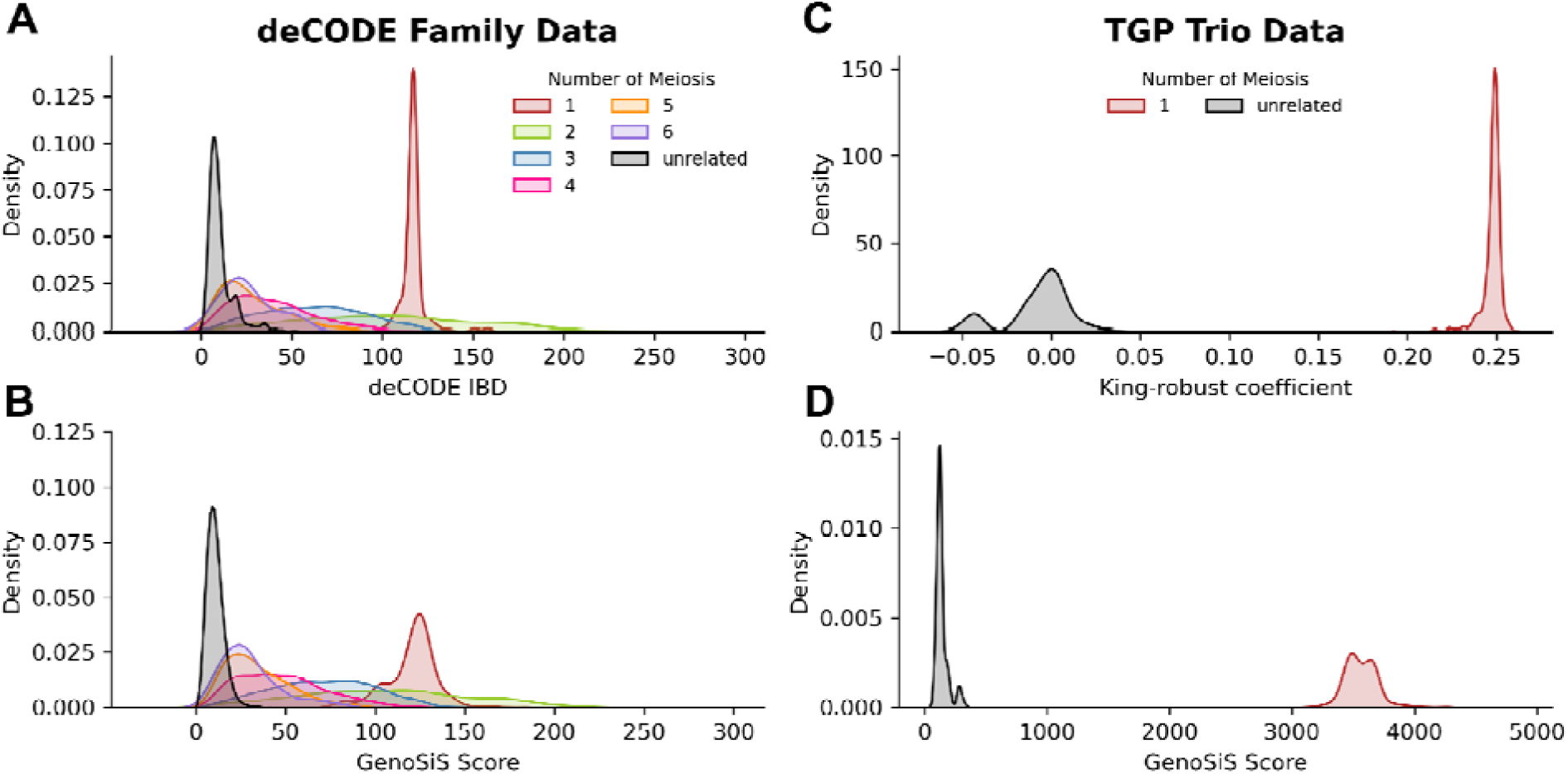
Genetic distance distributions among closely related samples. **A** deCODE IBD and **B** GenoSiS score distributions for deCODE families. **C** King-robust coefficient and **D** GenoSiS score distributions for TGP trios. The deCODE IDB and GenoSiS score ranges correspond to the number of cMs considered while the King-robust coefficient is normalized such that first-degree relatives have a score ∼0.25, second-degree relatives ∼0.125, and so forth. **AB** considered two haplotypes for chromosome 18. **CD** considered all autosomes. Distributions plotted and colored by numbers of meiotic events separating two samples where the number of meiotic events corresponds to family relationships (e.g. 1 = parent/child, 2 = grandparent/grandchild, full-siblings, etc.). All cohorts were size 20.

#### 1. Closely related individuals

We start by considering relatedness among biological family members because we have an expectation of how related two people are based on the number of meiotic events which separate them. On average, ∼50% of our autosomal DNA passes to our children, 25% to our grandchildren, and so on, but even at this scale, the randomness of recombination and other factors complicate these expectations. Full siblings can share less than 40% to more than 60% of their DNA and cousins can share between 5% and 20%^17,18^.

To analyze performance among closely related individuals, we used ten pedigrees (chromosome 18) from deCODE Genetics, each spanning four to six generations. For each sample, we generate GenoSiS cohorts where *k=20* and compare the GenoSiS’s scores (see Methods) between pairs of samples to their nominal relationship and their deCODE IBD metric (**Figure 2A,B**). Additionally, we compared the difference between the GenoSiS score and the King-robust kinship coefficient score among the 608 trios in the TGP cohort and between unrelated samples (**Figure 2C,D**).

In the deCODE genetics pedigrees and the TGP trios, the GenoSiS score showed a correlation with sample relatedness that was consistent with other methods. For ten deCODE pedigrees, each spanning four to six generations, the GeonoSiS score and deCODE’s IBD metrics easily distinguished between parent/child relationships and unrelated samples. While the distribution medians for more distant relationships decreased as the number of meiosis increased, as expected, there was significant overlap in the distributions, making reliable differentiation among these relationships difficult for either method (**Figure 2A,B**). Similarly, the trios (i.e., mother, father, child) in TGP were distinguished from unrelated samples using the Geonosis score and King-robust coefficient (**Figure 2C,D**). These results highlight the effectiveness of the GenoSiS score in quantifying genetic relatedness while also demonstrating the challenges inherent in differentiating more distant relationships.

#### 2. Distantly Related Individuals

While our expectations among more distantly related individuals are weaker—we are generally more related to those whose ancestors originated near our ancestors—this search is more relevant because biobank-derived cohorts are more likely to include non-family members. Given our relatedness expectations at this scale, we evaluated cohorts based on genetic ancestry specificity. Due to the complexity of human ancestry and population stratification, it is challenging to definitively state that a cohort should comprise *P*% of individuals from a specific group or that participants within a cohort should have a GenoSiS score of *S* or higher. For the analysis of distantly related samples, we can say that for a fixed cohort size (e.g. *20*), a cohort with more individuals from the same population as the query sample (i.e. 14/20 are from the same population as the query) is preferred over one with more individuals from a different population than the query sample (i.e. 14/20 are from a different population than the query). Further discussion about the quality of these cohorts depends on the population’s natural history, and some of this is addressed in section 4 and elaborated on in our discussion.

To evaluate cohorts formed from distantly related samples, we considered 3,202 TGP samples, which included 26 populations that we refer to as *subpopulations* (e.g., ESN=Esan, GBR=Great Britain, see Supplementary Table S1 for the full list), each of which falls into one of five super populations (AFR=African, AMR=American, EAS=East Asian, EUR=European, and SAS=South Asian). We retained all trios to mimic a real-world biobank where we may want closely related samples to appear in cohorts when available. We removed one sample which is listed as both AFR and EUR superpopulation. While the GenoSiS operates the same for sample of mixed ancestry, we wanted to keep our analysis for TGP data consistent for only samples with known, single ancestry. We used this data to compare GenoSiS to Plink’s DST and pi-hat metrics^6^ and the King-robust kinship coefficient estimator^7^. With GenoSiS, we queried individual samples and used their nearest neighbors to determine their unique custom cohorts for each individual. In contrast, the other three methods provide NxN pairwise relatedness scores. From these scoring matrices, we sorted the results for each sample and retained the top scores to form their custom cohort.

Cohorts drawn from a fixed population will inevitably include more diverse samples as they grow larger. To track the specificity of cohorts as they become more diverse, we sorted cohorts in decreasing relatedness to the query sample (i.e., a cohort with one sample includes only the most related sample and one with 20 includes the full cohort) and tracked the proportion of the cohort that matched the query’s super and subpopulation (**Figure 3A**). The TGP cohorts created by GenoSiS contained a higher density of samples matching the query at both the super and subpopulation levels. Between 98.97% and 100% of samples in the GenoSiS cohorts matched the query sample’s super-population, while the other three methods included up to 30% of samples from other super-populations. In nearly every instance, the GenoSiS cohorts retained more samples with matching subpopulations and never included more than 20% of different subpopulations. The one exception was the African population, where GenoSiS and the King-robust kinship coefficient retained subpopulation samples at about the same rate. See **Supplementary Figure S1** for comparison of GenoSiS with plink DST and pi-hat.

**Figure 3.**
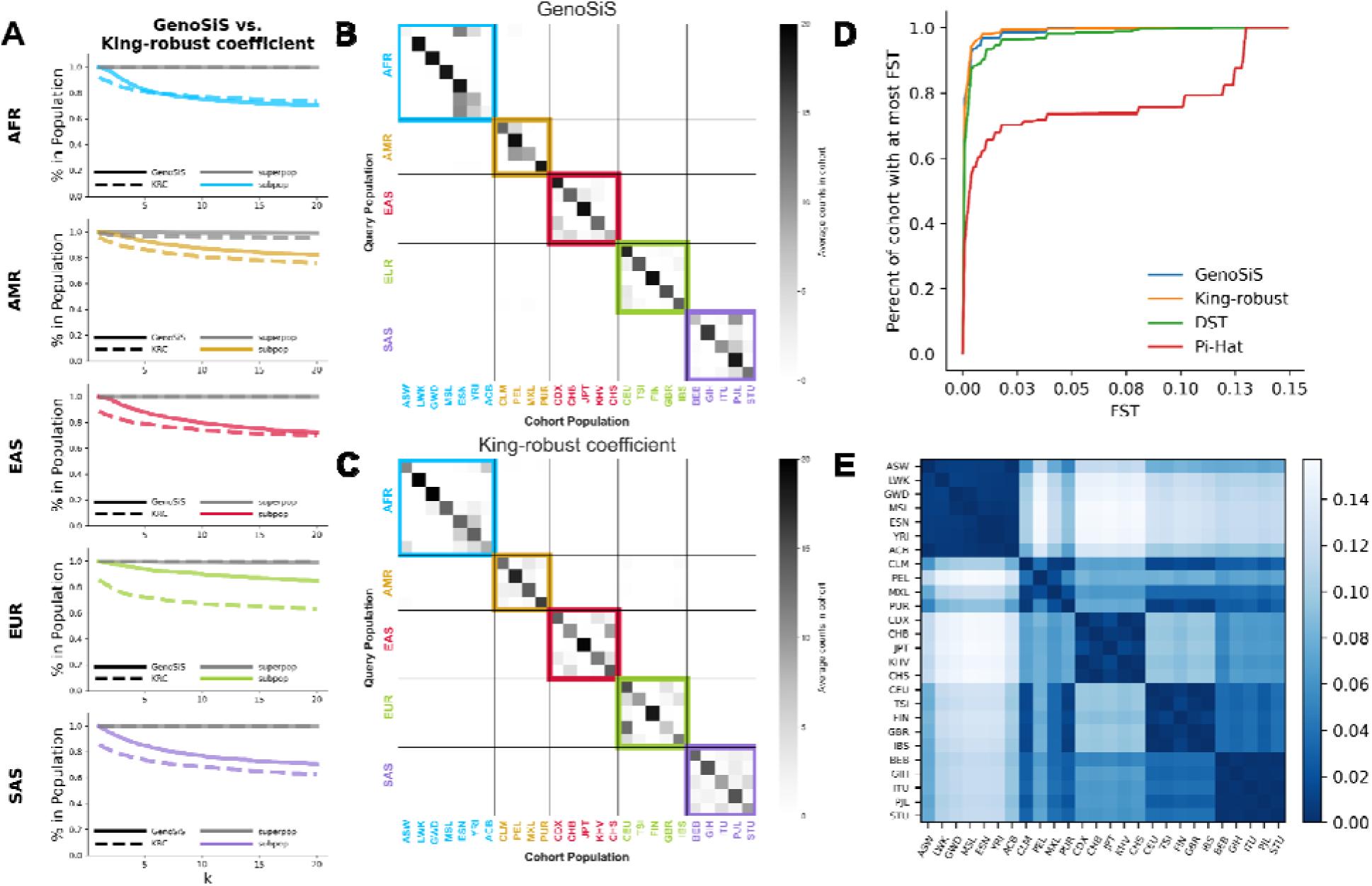
Analysis of TGP cohorts. **A** Percentage of samples from the TGP cohorts which are in the super and subpopulation of the query samples as the size of the cohorts increase from 1 to 20. GenoSiS is plotted as a solid lin while the King-robust coefficient is plotted as a dashed line. Lines for super populations are plotted in gray, while lines for subpopulation are plotted in a color corresponding to the population. Average subpopulation counts of appearances in cohorts generated by **B** GenoSiS and **C** King-robust coefficient. For both **B** and **C**, super population labels for query samples are listed on the vertical axis and are colored accordingly. Subpopulation labels for samples in the representative cohort are listed on the horizontal axis and are colored according to their respective super population. F_ST_ is a statistical measure of genetic differentiation between populations^21^. **D** Empirical distribution function tracking the percentage of samples identified by 3 comparison methods that were from subpopulations with an F_ST_ value less than equal to than a specified threshold. For instance, 78% of samples identified by GenoSiS were from populations with an F_ST_ ≤ 0.001, King-robust had 75.2%, DST had 64.8%, and pi-Hat had 34.6%. A lower F_ST_ and darker blue color indicates higher genetic similarity. **E** F_ST_ for all TGP subpopulations grouped by super population. Cohort size *k = 20* for all scenarios.

To understand the composition of the cohorts produced by each method at a more granular level, we organized the cohorts into groups based on the query sample’s subpopulation, and calculated the average occurrence of each subpopulation in each group (**Figure 3**). For example, the GenoSiS African Ancestry in Southwest US (ASW) cohorts (**Figure 3B**) had samples from three different super populations (95.6% African, 4% European, 0.4% American) and eight different subpopulations. Among the eight subpopulations, three were dominant: Esan in Nigeria (ESN), Yoruba in Ibadan, Nigeria (YRI), and African Ancestry in Southwest US (ASW), with an average of 11.3, 5.1, and 1.7 samples per cohort, respectively. This structure is consistent with our current understanding of African American genetic ancestry^19,20^.

GenoSiS produced cohorts with more in-subpopulation samples. Four of the 26 GenoSiS subpopulation cohorts contained exclusively samples matching the queries’ subpopulations, which did not occur with other methods. Most (65%) of the GenoSiS subpopulation cohorts contained at least 70% of samples matching the queries’ subpopulation. In comparison, only 19.2%, 34.6%, and 42.3% of cohorts produced by Pi-hat, DST, and King-robust kinship methods, respectively, met this criterion. Overall, GenoSiS and King-robust produced cohorts with samples that were from highly similar subpopulations based on subpopulation to subpopulation F_ST_ ^21^ (**Figure 3D,E**).

#### 3. Cohorts with poor representation

Identifying representative cohorts in a biobank relies on the biobank having individuals who are genetically similar to the patient. Because biobank diversity reflects regional diversity, health systems can expect to encounter patients who are poorly represented in their biobank. Therefore, it’s crucial to determine if a patient’s cohort is sufficiently representative.

To understand the dynamics of queries against underpowered databases, we used data from the TGP to construct six population-specific GenoSiS indexes and six population-specific query sets, then tracked the GenoSiS score for each query set against each index (**Figure 4**). As expected, GenoSiS scores were lower when the index and query set diverged. We can establish thresholds from these results to determine if a cohort adequately represents the query. For example, using TGP, we might exclude cohort members with a GenoSiS score below 1500 and assess whether the remaining members are sufficient for future analysis. While exact threshold will be specific to each database and type of analysis, these results clearly demonstrate the need to assess cohort representation.

**Figure 4.**
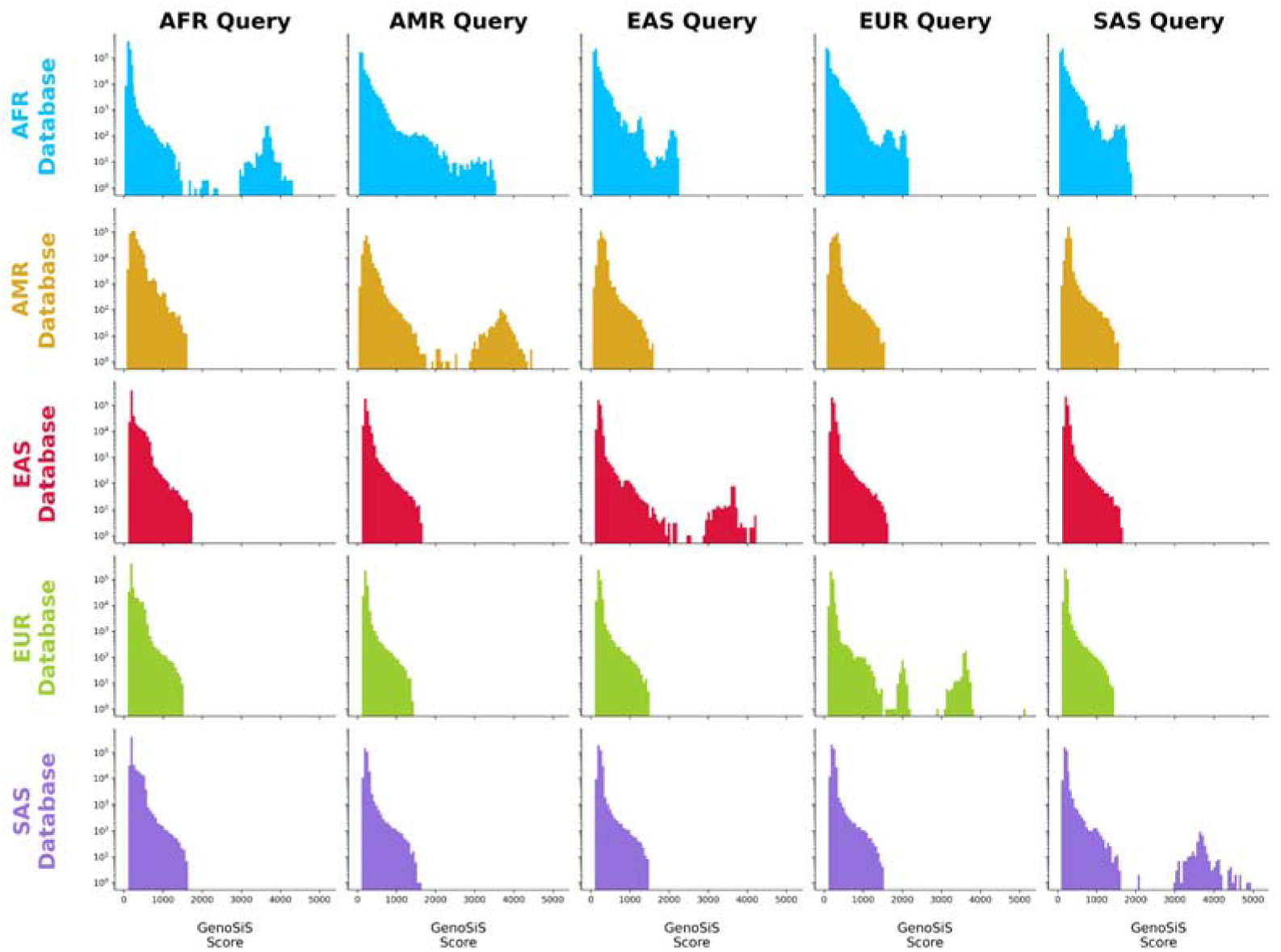
Cohort composition for queries with matched and mismatched TGP populations. Rows are representative and colored by the population of the database (i.e. samples which can appear in the GenoSiS cohort), columns are representative of the query population. In cases where the query and database sets were from the same super population (diagonal) we separately label when the cohort samples matched the query samples’ sub and super population labels.

#### 4. Potential Inequities

While GenoSiS performance is consistent across TGP populations, it is still critical to identify any potential inequities when training machine learning models. Here, we define an inequity as any instance where performance differs across populations. These inequities often arise from inadequate or biased training strategies and can typically be resolved by improving the training corpus. When improved training cannot resolve the bias, it must be detected, understood, and steps must be taken to mitigate its effects.

For example, in the embedding stage, GenoSiS converts a sample’s genotypes to a genetic embedding. Genotypes are then represented as two variable-length positional haplotype encodings with one entry for every non-reference allele, and two vectors per sample. Haplotype embeddings are a fixed length floating point vector. The conversion of a variable-length vector to a fixed-length vector has the potential to introduce bias if different populations have different haplotype vector lengths. The manifestation of this potential bias comes from a reference bias effect where individuals with African ancestry have more non-reference alleles and longer haplotype encodings (**Figure 5A**).

**Figure 5.**
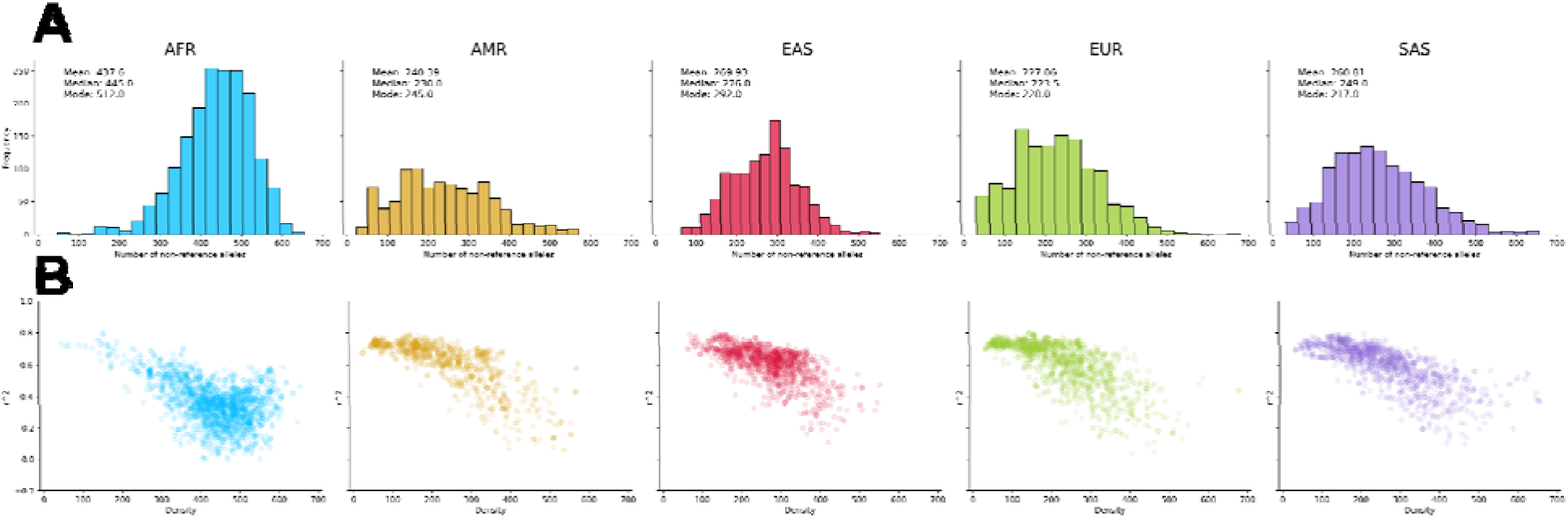
Embedding quality is worse for individuals with more non-reference alleles, who most often have African genetic ancestry. A The distribution of non-reference allele counts by population. B The relationship between a sample’s embedding quality, measured by the correlation coefficient (R²) between the genotype encoding and embedding distances, and the number of non-reference alleles in the sample’s genome.

To investigate this potential source of bias, we compared how the number of non-reference alleles in a sample corresponds to embedding quality. Our model learns embeddings by minimizing the difference between the distances of haplotype encoding pairs and the distances of the corresponding haplotype embedding pairs. We can measure the overall quality as the linear correlation (R²) between the genotype and embedding distances. Ideally, high correlations across populations would indicate no bias. However, the African population has a distinct decrease in R² (**Figure 5B**). Although this bias does not significantly affect our ability to find cohorts for this population, we plan to address the bias by developing a more robust training corpus that includes more examples with long genotype encodings. Modifying the training objective to penalize input encoding length could improve model performance without the need to expand the training set.

#### 5. Colorado Center for Personalized Medicine Biobank

The most compelling application of GenoSiS is in searching biobanks for cohorts genetically similar to patients. This search is similar to searching TGP, where the cohorts are mostly non-family members. However, unlike TGP, a biobank’s population structure is determined by its participants, often reflecting the regional diversity where the samples were collected. For example, the CCPM biobank’s^15^ diversity mirrors that of the broader Rocky Mountain region. Based on inferred genetic ancestry cluster memberships, the biobank consists of 80% European ancestry, 10% Hispanic individuals, 5% African American individuals, and 5% other underrepresented populations. The CCPM biobank is also significantly larger than TGP, with 73,346 samples, over 20X more samples than TGP.

Similar to our analysis for TGP populations, we evaluated GenoSiS cohorts at the population level using predicted genetic similarity groups to account for ambiguities in racial and ethnic categories or mis-classification of race/ethnicity during clinical visits. Ancestry inference was based on projections of CCPM samples to ancestry clusters in TPG and the Human Genome Diversity Project (HGP)^14^ reference panels. Samples are assigned a probability for each ancestry cluster and are then assigned the label of the cluster with the highest probability. CCPM adopts group labeling approaches which follow recommendations on population descriptors from the 2023 NASEM report^22^ and are defined at the super population level. See Supplementary Table S2 for CCPM labeling information. Given the size of the biobank, cohorts from other methods were not available for comparison.

GenoSiS found cohorts predominantly from populations that matched the query samples’ population (**Figure 6A**). The exceptions were from MLE- and SAS-like populations, where the cohorts were most similar to the trained categories of EUR-like ancestry. The discrepancies with the MLE- and SAS-like populations are driven by two factors likely to affect many biobank- backed clinical analyses: small populations and ambiguous ancestry prediction. The MLE- and SAS-like populations were least represented in the CCPM biobank, constituting just 0.3% and 1.6% of the total, respectively. Variable sample sizes are a known source of bias for methods that rely on dimensionality reduction^23–25^. Compounding the small populations was a shift in the quality of the ancestry inference for these two populations (**Figure 6B**). The MLE- and SAS-like cohorts also had the lowest-scoring cohorts (**Figure 6C**, **Supplementary Figure S3**). The issues arising from small populations and ancestry inference highlight the importance of genetic-distance-based cohort creation, which GenoSiS is the only method designed to accomplish at scale.

**Figure 6.**
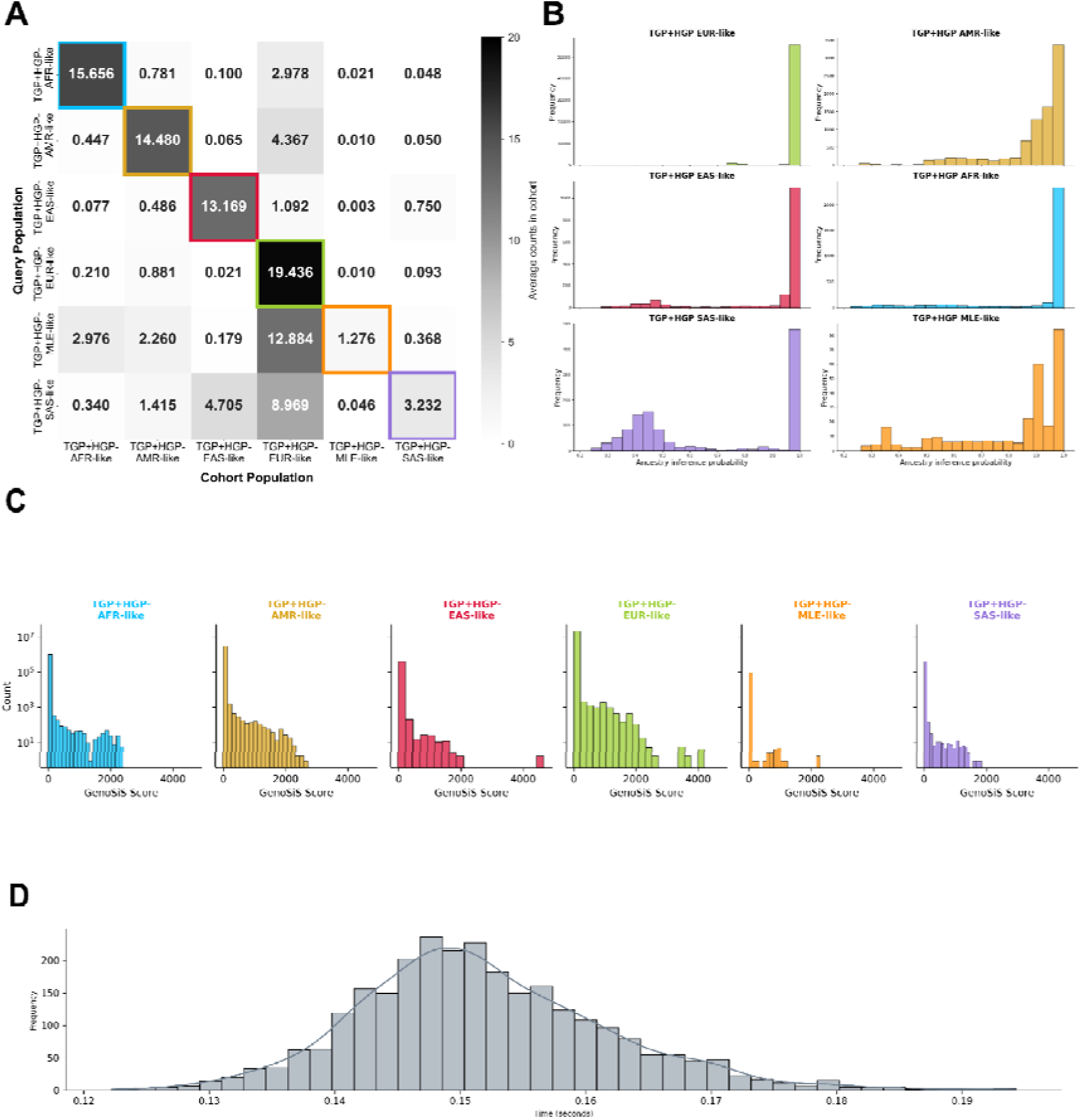
Average occurrence of each population for CCPM cohorts of size 20. **A** Average CCPM group counts for cohorts generated by GenoSiS plotted as a heatmap. Group labels for the query samples are listed on the left axis, while the labels of sample which appear in their GenoSiS cohort appear on the bottom axis. **B** CCPM ancestry inference probabilities for CCPM populations. **C** GenoSiS scores for CCPM groups. **D** GenoSiS search and aggregation times for CCPM’s 73,346 biobank samples.

### B. Query Runtime

As the number of samples in biobanks continues to grow and the demand for patient-centered results increases, the speed at which we can find cohorts will become critical to their usefulness. Among the 74,346 CCPM samples, GenoSiS can search for a single query in less than a tenth of a second (∼0.025 milliseconds) on a single c3-highmem-22 compute instance in Google Cloud. Since each query (one segment) can be performed independently (i.e., queries by segment do not depend on other segments), the runtime for a single sample (many segments) can be further improved using additional threads or specialized hardware accelerators. These results demonstrate that GenoSiS is fast enough to empower many patient-centric analysis, including point-of-care applications (**Figure 6D**).

## III. ONLINE METHODS

### A. Haplotype Embedding Model Training

#### Model architecture

The haplotype embedding model takes positional haplotype encoding vectors as input (see Methods B1) and outputs a 512-dimensional embedding vector. The model consists of a 1D convolutional neural network (CNN) with 6 blocks followed by a global mean pooling operation and a fully connected layer to output the embedding vector. Each convolutional block consists of a 1D convolution followed by dropout, batch normalization, *gelu* activation function, and max pooling. Each convolutional block has a kernel size of 6 and stride of 1. The number of input and output channels is defined by 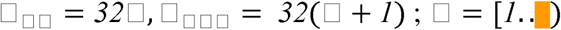 where i denotes the ith convolutional block. For the 0th block, we have 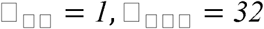 Max pooling kernel size is 3, and the output embedding dimension is 512.

#### Siamese Training

The CNN was trained in a Siamese fashion. During training, pairs of segment-matched positional haplotype encoding vectors were presented to two CNNs with shared weights. The training objective was to minimize the mean squared error between two output embeddings’ cosine similarity and the ground truth precomputed cosine similarity between their corresponding raw haplotype encoding vectors (see Methods B1). Gradients along both CNN paths were mean aggregated and then used to update weights. The model was trained for 200 *epochs* with a *batch size* of 1024. For the optimizer, we used *AdamW*^26^ with 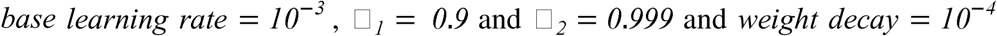. The base learning rate was scaled using torch’s *CosineAneallingWarmRestarts* learning rate schedule with parameters `T_0=17328`, `T_mult=1`, and `eta_min=1e-6.`

### Training Set

To train our model, we used positional encoding vectors (see Methods B1) from TGP data, chromosome 8 (169 segments, 3.8 million variants). As we are only interested in quantifying similarity between the same segments, we only train on pairs within the same segment. In this system, we choose cosine similarity as our ground truth and specify our output embedding vectors to be of length 512. See Supplementary Table S3 for the split of segments used for training, testing, and validation sets.

To obtain pairs of training samples that span a wide, uniform range of similarities, we do the following:

~~~
For each sample S in TGP:
    For each subpop P in TGP subpopulations:
        Randomly select 5 samples from P.
        Pair each random sample with S.
        Compute pairwise similarities for each pair.
Divide pairs by similarity values into 20 bins.
Randomly select at most 500 pairs from each bin.
~~~

We then split the 169 segments into train (80%), test (10%), and validation (10%) sets to ensure the model validates/tests on unseen genomic regions.

### B. The GenoSiS Architecture

The GenoSiS workflow consists of 4 major steps (Figure 1): **1) Data preprocessing**, where input VCF files are split into one centimorgan encoding segments, **2) generating embedding vectors** from segment encodings with a trained neural network, **3) building approximate nearest neighbors (ANN) search indices** from embedding vectors, and **4) searching the indices** to report the final cohort. A general overview can be seen in Figure 1, and further details for each step are described below.

#### 1. Data Preprocessing

##### Input and intermediate files

- VCF File: Genotype data should be formatted as a phased VCF file, compressed, and indexed with bgzip and tabix.
- Map file: Map files can be obtained from various sources, and serve as a reference for the interpolated map generated in an intermediate pre-processing step. We use the build GRCh38 PLINK-format genetic map out of Washington University, links provided below and in our GitHub repository.
- Interpolate map: To appropriately estimate start and end base-pair positions for the next step, we map an appropriate positional centimorgan to each base-pair coordinate that appears in the input VCF file. An intermediate map file is generated by interpolating positional centimorgan data for each base-pair position witnessed in the VCF using the centimorgan data provided in the reference map file.
- Segment boundaries: After the interpolated map file is created, the start and end points for all 1cM segments are recorded in a segment boundaries file.

##### Slice VCF

For most of the remaining steps (excluding the final aggregation step), we process 1cM segments independently. We choose a length of 1cM to align with the IBD-method consensus of minimum length for IBD (2-3cM). In this step, the input VCF file is parsed into 1cM segments starting and ending at the segment boundaries identified in the previous step. All output segments from this step are written to independent VCF files, including the original header. Once VCF slicing is complete, each procedure can be completed independently for each segment.

##### Binary haplotype encoding vectors

To generate binary haplotype encoding vectors, GenoSiS encodes non-reference and reference alleles as 1 and 0, respectively. GenoSiS encodes biallelic variants separately, creating two *binary haplotype encoding vectors* for each sample (one vector for each of two genetic parents, or haplotypes). First, we record all genotypes in a data structure we call “variant major format (VMF),” where rows are variants and columns are samples (as formatted in a VCF file). From VMF, we generate transpose to sample major format (SMF), where rows are samples, and columns are variants.

##### Positional haplotype encoding vectors

Binary haplotype encoding vectors are extremely sparse. From these sparse encodings, we create dense, positional haplotype encoding vectors. Here, we encode the centimorgan position (from the interpolated map file) only at sites marked as non-reference. By encoding the centimorgan position of a non-reference allele, we preserve the location of variants along a segment.

#### 2. Generate embedding vectors

With our trained model (See Methods A), we generate *haplotype segment embedding vectors* for all segments in all chromosomes by running inference on the model using the *haplotype positional encoding vectors* as input.

#### 3. Creating Index

##### SVS Index

For each 1cM segment’s embedding vectors, we build an ANN search index using default settings from the Intel® Scalable Vector Search (SVS). While SVS is one of the highest performing vector search databases, there are other solutions^27^ that could be used in its place.

#### 4. Searching Index

##### SVS search

A search takes haplotype segment embedding vectors, and SVS segment indices as input and returns the k-nearest neighbors for each query sample at each segment. Here, the selection of *k* is a tunable parameter, and we consider the significance and interpretation of different *k*-values in our discussion section.

#### 5. Aggregate segments

To report a genome-wide representative cohort, we aggregate SVS search results from each segment’s index into a binary vector denoting presence or absence in top *k* for all segments in a single chromosome. Samples with high similarity to the query will be dense, and those with low similarity will be sparse. To report a final cohort, we perform a summation operation for all match vectors. For aggregation at the chromosome level, we add up sample scores for each chromosome and report a single score for all match-sample IDs that appeared one or more times in the top *k* across all segments. The final cohort considers all match scores from each chromosome, ranks them by order of decreasing values, and reports top *k* samples, genome-wide.

### C. Experiments

#### 1. Closely related individuals

Tracking relatedness among family members by number meiosis is complicated because higher degrees of relatedness include various relationships with different inheritance patterns. For example, degree 2 includes both grandparent/grandchild and full siblings. Full siblings share genetic data on two haplotypes while grandparent/grandchild share genetic data only on one haplotype. This arrangement leads to varying amounts of shared DNA beyond straight forward percentage expectations. This pattern continues with increased degrees of relatedness to include more and more relatives.

Using six families from deCODE’s database and 608 trio samples from TGP, we evaluate the performance of GenoSiS on closely related individuals. For the deCODE analysis, we create a GenoSiS index for the complete set of all families and query all samples to generate their GenoSiS cohorts. Similarly, for the TGP trio analysis we created a single index for all 608 samples and queried all samples for their GenoSiS cohort. We label pairs of individuals by the number of meiotic events (i.e. generations) by which they are separated, and add a label ‘unrelated’ for individuals who come from separate families. We benchmarked the correctness of deCODE GenoSiS cohorts against an IBD tool provided by deCODE. This tool, like other IBD methods, computes pairwise IBD for all samples to quantify length of shared segments of DNA between two samples. deCODE’s IBD method, like most IBD methods, does not report a genome-wide score, instead it reports pairwise IBD segments. The score reported in Figure 2a is a summation of IBD segments across both haplotypes.

#### 2. Distantly related individuals

Using TGP data with labels for super (e.g. AFR=African, EUR=European, etc.) and subpopulation (e.g. ESN=Esan, GBR=Great Britain) ancestry groups, we further investigate GenoSiS at the population level. We create a GenoSiS index for the complete TGP dataset and query all samples to generate their GenoSiS cohorts. For each query, we record the subpopulation label of the query, and of the *k* cohort members (e.g. a query sample who is South Asian (SAS), Gujarati (GIH) might have a cohort with 15 SAS-GIH, 3 SAS-BEB, 1 SAS-ITU, and 1 EUR-TSI). In Figure 3A, we demonstrate the decay of GenoSiS cohorts as more samples are included in the cohort (i.e. k increases from 1 to 20), and compare to cohorts inferred by other methods. We also report the average counts for all queries’ cohort populations and plot a heatmap of these averages in Figure 3B,C. We did the same for plink DST, pi-hat, and kinship and plot their results in Supplementary Figure S1. As discussed, none of the plink tools report a representative cohort (i.e. KNN) in the same way GenoSiS does. To generate plink cohorts, we added a post-hoc ranking step which selects the top *k*=20 samples with the best plink scores (i.e. most similar samples) for each query sample.

#### 3. F_ST_

**F_ST_** values for TGP subpopulation pairs were obtained from^21^. In Figure 3D. TGP subpopulations are grouped on each axis such that they are adjacent to their respective superpopulations, then we generate a simple heatmap of □_□□_ (□, □), for all combinations of subpopulations (□, □). In Figure 3E, for each TGP subpopulation, we use GenoSiS and other tested methods to aggregate cohorts using previously described methods and count the number of results from each subpopulation. We then plot the percentage of samples in each cohort from subpopulations with an F_ST_ value less than or equal to □ where □ ϵ [*0, 0.15*] to generate an empirical density function for each method.

#### 4. CCPM

We repeat the same process we used to generate Figure 3B,C to generate Figure 4A,B using more than 73K samples from the CCPM biobank. As there are no subpopulations associated with this dataset, we label only CCPM group labels. We compute **F_ST_** values for all 22 autosomes from CCPM data with Plink2’s --fst option.

#### 5. Quality

To investigate how GenoSiS performs on different database/query combinations, we generated 5 separate databases for each of the five TGP super populations (AFR, AMR, EAS, EUR, SAS). We then performed queries for all samples in all combinations of database/query combinations (e.g. all AFR sample query into an AFR database, all AFR samples query into an EAS database, etc.) In these experiments there are 895 African samples, 491 American samples, 586 East Asian samples, 634 European samples, and 602 South Asian samples. Figure 5 shows the results of these queries as a histogram of GenoSiS scores.

#### 6. Timing

To evaluate the speed of GenoSiS, we report timing for our largest available dataset (CCPM Biobank data). In compliance with CCPM data sharing agreement, we perform our search on a c3-highmem-22 compute instance in Google Cloud (176 GB, Intel Xeon Platinum 8481C @ 2.70 GHz) provided to us by our Health Data Compass support team. We report search times for a single query–where each sample constitutes two queries (i.e. two haplotypes)–over all 2,737 segments from the CCPM data. Each segment’s search time was reported individually, and the distribution of these times are shown in Figure 4D.

## V. DISCUSSION

GenoSiS is a rapid, genome-wide search for genetically similar individuals which scales to large biobanks. GenoSiS is inspired by early genomic similarity detection methods while leveraging new technologies to rapidly match query samples to representative cohorts. With GenoSiS, we move toward inclusive research in precision medicine and motivate further research for query-based cohorts over finite ancestry labels. Rapid and accurate identification of these representative cohorts can be used to support real-time clinical studies and develop evidence-based precision treatment plans for physicians, researchers, and other members of clinical teams. Additionally, the identification of these cohorts could have promising implications in the exciting pursuit of inclusive Genome Wide Association Studies (GWAS), polygenic risk scores (PGS), and other statistical analyses as biobank diversity continues to catch-up with our increasingly diverse global human population^28^.

A key risk to finding patient-matched cohorts is when insufficient patient diversity within an institution’s biobank fails to create a large enough cohort (see Section II.4). We can greatly reduce this risk by expanding the pool of potential patient matches via other institutions’ biobanks, so long as patient data remains private. One solution is a federated machine learning approach^29^, where each institution trains a standardized learning model on its own local data and shares only the learned model parameter updates with a central server for aggregation, a strategy that precludes the need to share raw data. We believe that GenoSiS can easily accommodate this model by adopting a common training set curation process across collaborating institutions and by leveraging ideas from standard federated learning for handling heterogeneous data. In this setting, GenoSiS will need to work across private databases housing different subpopulations, some or all of which may not share statistical distributions with the queries. The recent introduction of a form of dimensionality reduction into SVS [https://openreview.net/forum?id=wczqrpOrIc] that improves search recall across differing subpopulations paves the way for GenoSiS to be applied in a federated setting.

### GenoSiS performance on deCODE closely related individuals

Treating deCODE’s IBD method as a reliable measure of truth, Figure 2A supports the claim that relatedness for family members up to 6 degrees is complicated. While there is a strong signal for samples that are separated by 1 meiotic event (e.g. parent/child) and samples which are unrelated, the remaining signals (e.g. grandparent/child siblings, cousins, etc.) are less defined. GenoSiS is not an IBD method, but by designing the search to perform at the 1cM segment level allows for a reasonable comparison to these IBD results. Figure 2B demonstrates that GenoSiS scores have a similar distribution to IBD scores, with small differences. We note that GenoSiS scores for degree 1 relatives have a wider distribution than IBD scores, and GenoSiS scores are slightly right-shifted. Though the distributions are mostly in agreement, the discrepancies are likely explained because IBD methods prefer to be careful when labeling IBD segments. Many IBD methods, for example, will only label regions IBD if the segment is ≥3 cM in length. The current aggregation and scoring method for GenoSiS is not designed to consider this level of granularity. Furthermore, when scores exceed the ∼50% shared cM (e.g. pairs separated by 2 meiotic events), we note these samples are siblings and share genetic data on more than one haplotype.

### GenoSiS performance on TGP ancestry populations

Much like relatedness, uncovering population structure for human genetic data is a complex task. As discussed in our introduction, we elect to examine population structure from the lens of ancestry, where ancestry labels are assigned within the TGP data. The assumption of truth for these analyses (Figure 3) is that a cohort with more samples labeled with the same subpopulation as the query is more correct than a cohort with samples labeled with the same super population as the query is more correct than a cohort with samples labeled outside of the query’s super population.

To evaluate GenoSiS cohort correctness against this standard, we benchmark against cohorts defined by 3 plink scores: (1) DST, (2) pi-hat, and (3) kinship. Plink’s DST score is a rough approximation for pairwise IBS distance, where IBS distance is a function of measuring the number of markers at IBS0 (e.g. AA, aa), IBS1 (e.g. AA, Aa), and IBS2 (Aa, Aa). DST values range from 0 to 1. A score of 0 indicates no genetic relatedness while a score of 1 indicates genetic clones. Plink’s pi-hat score is a measure of proportion IBD. Likewise, a pi-hat score of 0 indicates no genetic relatedness while a score of 1 indicates genetic twins. Finally, the King-robust estimator performs relationship inference to estimate pairwise kinship coefficients. Here, duplicate samples have kinship scores of 0.5 (not 1), first-degree relatives have scores ∼0.25, second-degree relatives have scores of ∼0.125, and so on. Furthermore, negative kinship scores indicate samples from different populations. The distribution of GenoSiS and plink scores is shown Supplementary Figure S2.

Figure 3 demonstrates cohort correctness at the subpopulation level. First, we note that for all four analyses there is a strong diagonal signal for super population labels (outlined in colored boxes), indicating that each method can at least identify cohorts which mostly match a query’s super population. Second, in all four analyses there is a slightly weaker, but still apparent diagonal signal for most subpopulations, indicating that these methods often identify cohorts which largely match a query’s subpopulation. Evaluation at the subpopulation level demonstrates some unique patterns between these methods. For example, plink pi-hat (Supplementary Figure S1B) shows a strong signal where samples of Great British (GBR) ancestry appear in cohorts for query samples which are not of GBR (nor European) ancestry. This pattern is particularly noticeable for query samples of African ancestry. Plink DST cohorts (Supplementary Figure S1A) identify cohorts more consistent with the query subpopulation, but still occasionally include samples outside of a query’s super population (e.g. African and American queries). King-robust coefficient and GenoSiS cohorts consistently generate the most correct cohorts at the super and subpopulation level, with GenoSiS cohorts being slightly more in line with subpopulation labels than King-robust coefficient.

Figure 3B,C supports results shown in Figure 3A with decaying values of *k*. All methods perform well for most super populations (gray lines), but the subpopulation level shows some performance differences. For each method, GenoSiS reports equal or better cohorts as k values increase from 1 to 20.

### GenoSiS performance on CCPM data

Analysis on the CCPM Biobank samples, shown in Figure 4, demonstrates a similar pattern to the analysis on TGP samples. That is, for most ancestry groups, GenoSiS cohorts comprise other samples from the same ancestry as the query sample. With CCPM data, however, GenoSiS cohorts for samples of TGP+HGP-MLE-like and TGP+HGP-SAS-like ancestry comprise a higher proportion of TGP+HGP-EUR-like samples on average than would be expected. Supplementary Figure S3 further reports GenoSiS scores for the same set of data. This analysis shows that while there is an overrepresentation of European samples in queries for TGP+HGP- MLE-like and TGP+HGP-SAS-like samples, the GenoSiS scores for these TGP+HGP-EUR-like samples remain relatively low (i.e. not inflated over the GenoSiS scores for samples of the same ancestry group). Furthermore, some of the poorer performance of TGP+HGP-MLE-like and TGP+HGP-SAS-like populations could be due to how population groups were defined in CCPM. During this process, every individual is assigned to a single group even if there is nearly the same amount of support for assignment to a different group (e.g. admixed samples), which could convolute some of these group assignments. Of the 73,346 samples in the CCPM Biobank, about 2% are TGP+HGP-EAS-like, 1.6% are TGP+HGP-SAS-like, 4.1% are TGP+HGP-AFR-like, 0.3% are TGP+HGP-MLE-like, 80% are TGP+HGP-EUR-like and 12% are TGP+HGP-AMR-like. We note that the two ancestry groups with the poorest cohort quality (higher number of out-population samples in the cohort) also have the fewest number of samples represented in the database. It has been well established that an imbalanced set of classes (i.e. ancestry groups) complicates knn search algorithms^30,31^. The consequence nor the solution for this imbalance is straightforward in this context, but we investigate the limitations of these imbalances and other possible biases, and discuss below.

### GenoSiS performance with different database-query representations

As shown in Figure 3B,C, and discussed above, the quality of a GenoSiS cohort can depend on the composition of the database of samples. For example, if GenoSiS creates an index on a database of exclusively European individuals, the GenoSiS cohort for an African query sample will necessarily constitute only European individuals, likely with low GenoSiS scores. To further investigate the scores for GenoSiS cohorts generated in these scenarios, we created five isolated databases for each of the five super populations in the TGP dataset and performed isolated queries for each pair of super population groups. The results from these experiments are shown in Figure 5.

As expected, we first observe that for all query populations the databases which return the highest GenoSiS scores are those which have the same population as the query. Further, we observe that the samples which contribute most to these higher scores come from samples within the same *subpopulation* as the query sample. A second observation is that for all queries, the African database returns higher GenoSiS scores for non-African queries than other databases do for out population queries. Specifically, queries of American ancestry observe cohorts with higher GenoSiS scores for African databases than for other databases. This observation is consistent with what we know about TGP American samples having admixed ancestry and TGP African samples having a more diverse representation over other TGP populations^19,20^. We include additional comments regarding the task of better understanding GenoSiS cohort quality for real-world databases (i.e. biobanks), and other comments on the current limitations of GenoSiS in our section on future work.

### Future work and limitations

At each step we offer comments about areas of current limitations and future investigation for the GenoSiS pipleline.

#### Data Preprocessing

Future versions of GenoSiS could benefit from analysis of optimal slice sizes (e.g. > 1 cM, < 1cM, variable slice sizes, etc.).

#### Encoding Vectors

GenoSiS generates positional encodings derived from the binary encodings, however, nominal encodings should be considered in place of binary encodings to account for the multiallelic nature of human variation.

#### Siamese Training

Future work should focus on serving underrepresented groups by asserting a more robust training process. Some possibilities might be to include more admixed populations and high variation samples, exploring the option to select a more representative chromosome or set of segments across many chromosomes, adding features of the data (e.g. multiallelic sites, structural variants, etc.), adding other tasks (e.g. maximize differences between ancestry groups), or training on different distance metrics, especially if nominal encoding vectors are added. Applying Summix^32^–a tool which provides ancestry adjusted allele frequencies–to training data would be an immediately useful step to address more than one of these ideas.

Furthermore, if other types of data are considered as input to the model (e.g. methylation, expression, health records, etc.), new distance metrics could be considered. The architecture of a Siamese training process allows for any distance metrics to be applied in the loss function; and downstream steps of GenoSiS are already designed to easily handle different kinds of embedding vectors. As such, the applications for new data and more complex training decisions are reasonable additions.

#### SVS index and search

More specific recommendations for choosing an appropriate k should be quantified. Currently, GenoSiS is set up to run with mostly default options, using the Vamana index with L2 distance. Future work should explore other SVS indexing, searching and loading options. Advanced vector compression techniques in SVS [https://dl.acm.org/doi/abs/10.14778/3611479.3611537] will provide further acceleration and memory footprint reductions.

#### Aggregation

During development, we investigated other methods for aggregation including merging gaps, setting a minimum limit to consecutive segment inclusion, and using the reported SVS score as a scoring technique. Popcount was most consistent with our initial evaluation criteria, but assert that further exploration of these and other scoring algorithms (e.g. HMM) have interesting paths forward.

#### Application

Applying GenoSiS

## Acknowledgements

This research was supported by NIH 1R01HG011774-01A1. Colorado Center for Personalized Medicine: John Finigan, Nicholas Rafaels, Health Data Compass: Gabriel Labrie, Michael Carter, Anna Lee, Matthew Fisher. Intel Labs: Ted Willke, Mark Hildebrand. BioFrontiers IT: Matthew Hynes-Grace. Sequence analysis lab at deCODE genetics, specifically Marteinn Hardarson and Bjarni Halldorsson. Regeneron Pharmaceuticals, Inc.

## Author Credit

**Kristen Schneider**–conceptualization, methodology and software development, data analysis and experimentation, manuscript writing. **Murad Chowdhury**–neural network software development and training, manuscript review and editing. **Mariano Tepper**–SVS software development and support, manuscript review and editing. **Colorado Center for Personalized Medicine**–CCPM Data curation, storage, and distribution. **Jawad Khan**–SVS support, supervision, manuscript review and editing. **Jonathan Short**–CCPM data curation and analysis support, manuscript review and editing. **Chris Gignoux**–CCPM data curation, analysis support and collaboration supervision, manuscript review and editing. **Ryan Layer**–supervision, conceptualization, data curation, collaboration supervision, data analysis and experimentation, manuscript writing, review and editing.

## Data and Code availability

### Thousand Genomes Data (TGP)

We use phased VCFs for 22 autosomes from TGP 30x on GRCh38. 3,201 samples are labeled 5 “super populations” (African, American, East Asian, European, and South Asian) and 26 “subpopulations”. One sample with AFR-EUR descent is omitted from analyses.

### deCODE families

From the deCODE database, we isolated chromosome 18 for ten families– each which had a least four and up to six generations–totaling 274 samples. Siblings which were not full-siblings were pruned from the dataset.

### Colorado Center for Personalized Medicine (CCPM)

We use phased VCFs for 22 autosomes from the CCPM database. Of the 73,346 samples, roughly 23k were genotyped with the Mutli-Ethnic Genotyping Array, while the remaining ∼50k had exome sequences with a genomic backbone. All samples were imputed with Topmed version 2. Ancestry labels are as follows: 1,445 TGP+HGP-EAS-like, 1201 TGP+HGP-SAS-like, 3041 TGP+HGP-AFR-like, 254 TGP+HGP-MLE-like, 58708 TGP+HGP-EUR-like, and 8697 TGP+HGP-AMR-like.

### Map files

https://bochet.gcc.biostat.washington.edu/beagle/genetic_maps/

In addition to the software to run GenoSiS, we provide helper scripts to move between some common file formats and generate necessary input.

Software available for download at GitHub: https://github.com/kristen-schneider/precision-medicine.

Figures and analysis scripts available at GitHub: https://github.com/ryanlayerlab/genosis_analysis

## Supplementary Information

**Figure S1.**
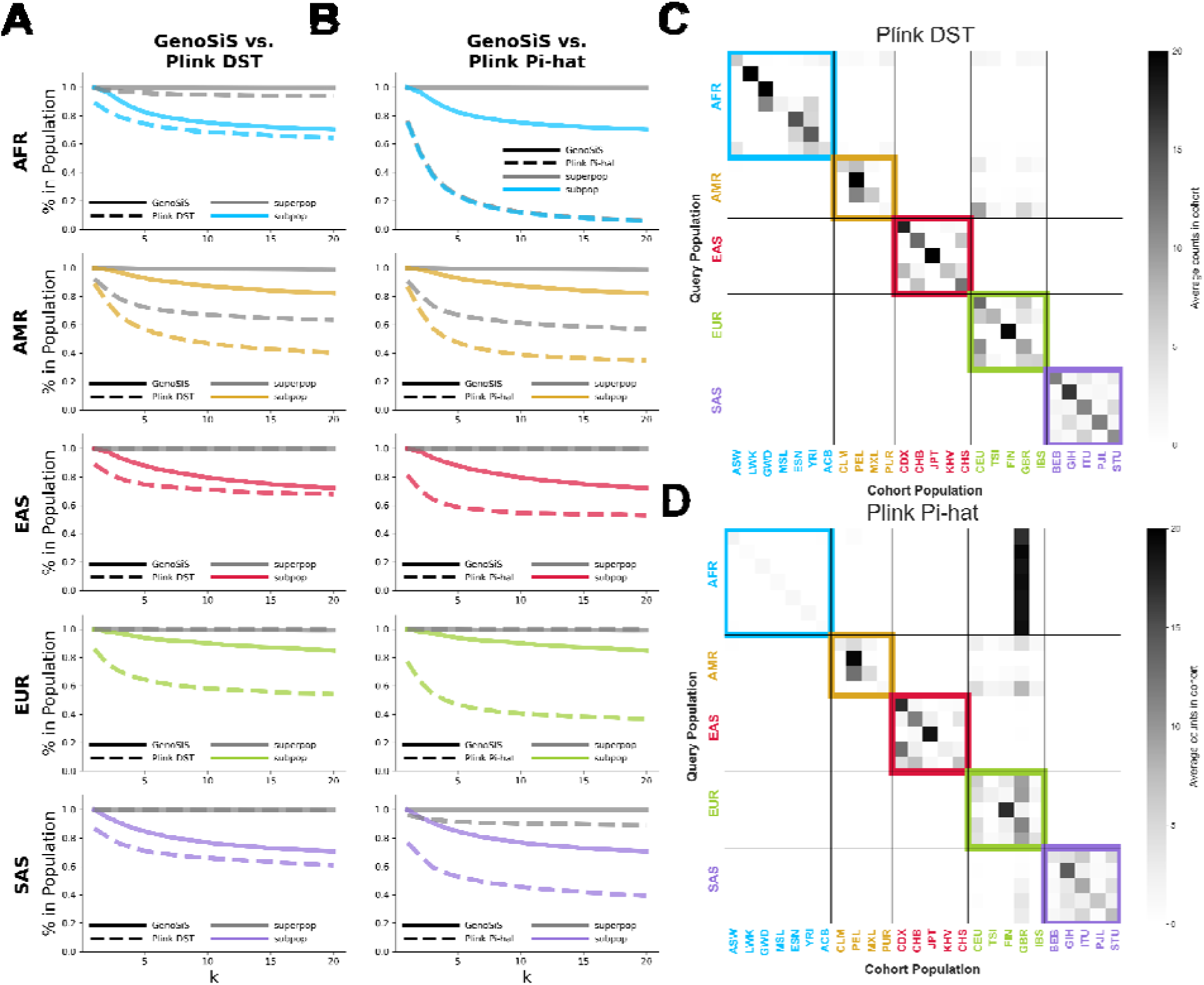
Super- and subpopulation percentages for TGP cohorts. **AB** The percentage of samples from the cohort which are in the super and subpopulation of the query samples as the size of the cohorts increase from 1 to 20. GenoSiS is plotted as a solid line while Plink DST **(A)** and Plink Pi-hat **(B)** is plotted as a dashed line. Lines for super populations are plotted in gray, while lines for subpopulation are plotted in a color corresponding to the population. **C** Plink DSK and **D** Plink Pi-hat. For both **C** and **D** super population labels for query samples are listed on the vertical axis and are colored accordingly. Subpopulation labels for samples in the representative cohort are listed on the horizontal axis and are colored according to their respective super population.

**Figure S2.**
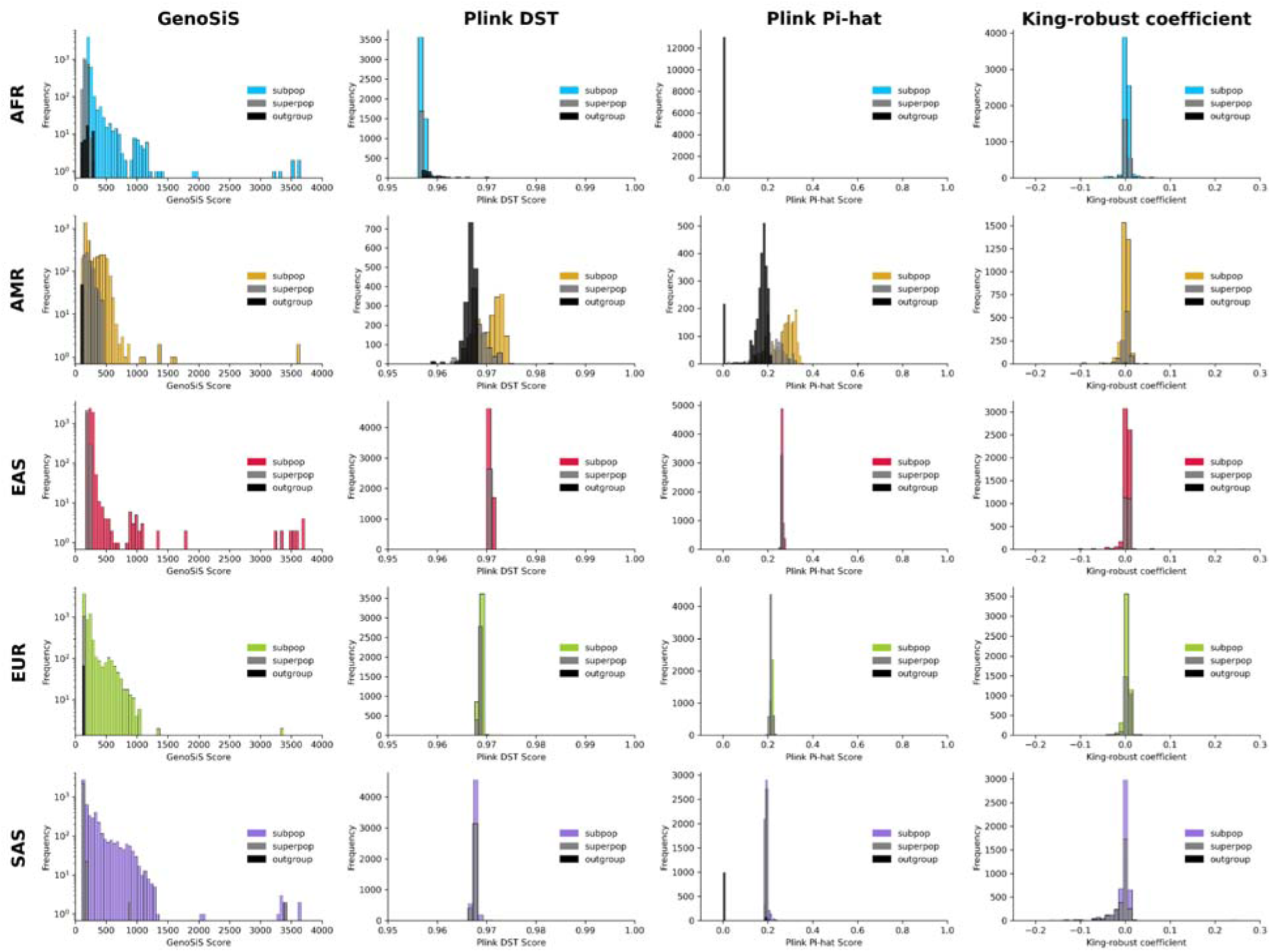
TGP cohort scores by population. Histograms of GenoSiS, plink DST, plink pi-hat, and King robust coefficient cohort scores for TGP data for *k*=20. TGP super populations are organized by row. Samples in the cohor which appear in the same subpopulation as the query sample are colored in yellow, red, green, blue, and purple for AFR, AMR, EAS, and SAS query samples accordingly. Samples in the cohort which appear in the same super population as the query sample are colored in gray. Samples in the cohort which appear outside of the query’s super population are colored in black.

**Figure S3.**
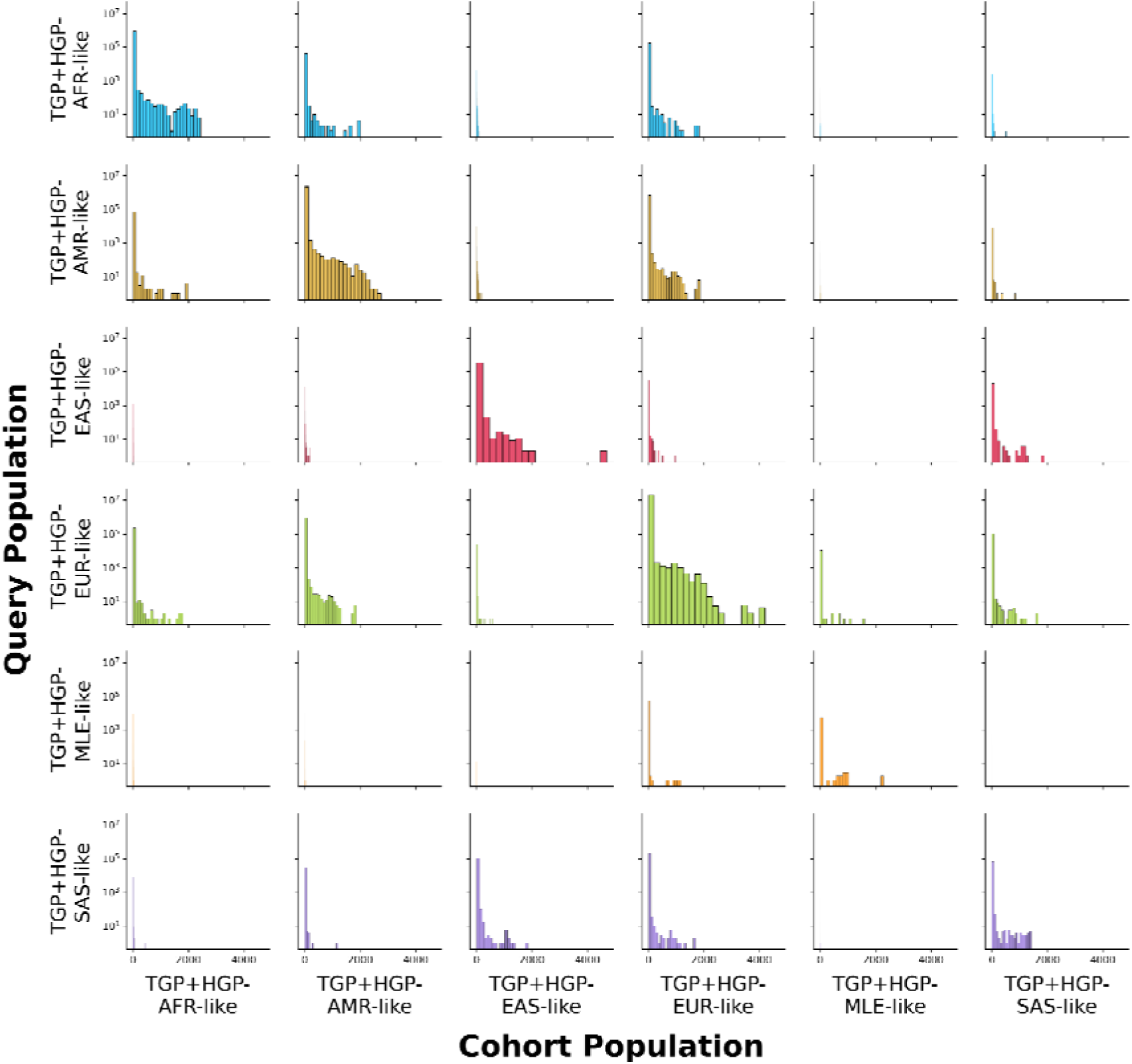
CCPM Biobank cohort scores by population. Histograms of GenoSiS cohort scores when k=20 for CCPM Biobank data. Query and cohort population labels are plotted in the same order for both axes.

**Supplementary Table S1.** Thousand Genomes Project (TGP) Subpopulation and super population labeling.

| **Super Population** | **Subpopulation Abbreviation** | **Subpopulation Name** |
| --- | --- | --- |
| African (AFR) | ACB | African Caribbean in Barbados |
| African (AFR) | ASW | Americans of African Ancestry in SW USA |
| African (AFR) | ESN | Esan in Nigeria |
| African (AFR) | GWD | Gambian in Western Divisions in the Gambia |
| African (AFR) | LWK | Luhya in Webuye, Kenya |
| African (AFR) | MSL | Mende in Sierra Leone |
| African (AFR) | YRI | Yoruba in Ibadan, Nigeria |
| American (AMR) | CLM | Colombians from Medellin, Colombia |
| American (AMR) | MXL | Mexican Ancestry from Los Angeles, USA |
| American (AMR) | PEL | Peruvians from Lima, Peru |
| American (AMR) | PUR | Puerto Ricans from Puerto Rico |
| East Asian (EAS) | CDX | Chinese Dai in Xishuangbanna, China |
| East Asian (EAS) | CHB | Han Chinese in Beijing, China |
| East Asian (EAS) | CHS | Southern Han Chinese |
| East Asian (EAS) | JPT | Japanese in Tokyo, Japan |
| East Asian (EAS) | KHV | Kinh in Ho Chi Minh City, Vietnam |
| European (EUR) | CEU | Utah Residents (CEPH) with Northern and Western European Ancestry |
| European (EUR) | FIN | Finnish in Finland |
| European (EUR) | GBR | British in England and Scotland |
| European (EUR) | IBS | Iberian Population in Spain |
| European (EUR) | TSI | Toscani in Italy |
| South Asian (SAS) | BEB | Bengali in Bangladesh |
| South Asian (SAS) | GIH | Gujarati Indian in Houston, TX |
| South Asian (SAS) | ITU | Indian Telugu in the UK |
| South Asian (SAS) | PJL | Punjabi in Lahore, Pakistan |
| South Asian (SAS) | STU | Sri Lankan Tamil in the UK |

**Supplementary Table S2.** Colorado Center for Personalized Medicine (CCPM) ancestry group-labeling.

| Abbreviation | Full CCPM labeling |
| --- | --- |
| TGP+HGP-AFR-like | Thousand Genomes Project + Human Genomes Project-African-like |
| TGP+HGP-AMR-like | Thousand Genomes Project + Human Genomes Project-American-like |
| TGP+HGP-EAS-like | Thousand Genomes Project + Human Genomes Project-East Asian-like |
| TGP+HGP-EUR-like | Thousand Genomes Project + Human Genomes Project-European-like |
| TGP+HGP-MLE-like | Thousand Genomes Project + Human Genomes Project-Middle East-like |
| TGP+HGP-SAS-like | Thousand Genomes Project + Human Genomes Project-Central South Asian-like |

**Supplementary Table S3.** Embedding vector generation, segment split for training, testing, and validating steps.

| Genotype Embedding Model Process | Segments (Chromosome 8) |
| --- | --- |
| Training | 0, 1, 2, 3, 4, 5, 6, 8, 9, 11, 12, 13, 15, 16, 17, 19, 21, 22, 23, 27, 28, 30, 31, 32, 33, 34, 35, 36, 37, 38, 39, 40, 41, 42, 43, 44, 45, 46, 47, 49, 50, 51, 52, 53, 54, 55, 56, 57, 59, 61, 63, 64, 65, 67, 68, 69, 70, 71, 73, 75, 76, 77 |
| Testing | 14, 18, 20, 26, 29, 62, 72, 74 |
| Validating | 7, 10, 24, 25, 48, 58, 60, 66 |

## References

1. Ding, Y. et al. Polygenic scoring accuracy varies across the genetic ancestry continuum. Nature (2023) doi:10.1038/s41586-023-06079-4.

2. Aguerrebere, C., Bhati, I., Hildebrand, M., Tepper, M. & Willke, T. Similarity search in the blink of an eye with compressed indices. (2023).

3. Danecek, P. et al. The variant call format and VCFtools. Bioinformatics 27, 2156–2158 (2011).

4. Browning, B. L., Tian, X., Zhou, Y. & Browning, S. R. Fast two-stage phasing of large-scale sequence data. Am. J. Hum. Genet. 108, 1880–1890 (2021).

5. Bromley, J., Guyon, I., LeCun, Y., Säckinger, E. & Shah, R. Signature verification using a‘ siamese’ time delay neural network. Adv. Neural Inf. Process. Syst. 6, (1993).

6. Purcell, S. et al. PLINK: A Tool Set for Whole-Genome Association and Population-Based Linkage Analyses. Am. J. Hum. Genet. 81, 559–575 (09/2007).

7. Manichaikul, A. et al. Robust relationship inference in genome-wide association studies. Bioinformatics 26, 2867–2873 (2010).

8. McVean, G. A genealogical interpretation of principal components analysis. PLoS Genet. 5, e1000686 (2009).

9. Nicholson, G. et al. Assessing population differentiation and isolation from single-nucleotide polymorphism data. J. R. Stat. Soc. Series B Stat. Methodol. 64, 695–715 (2002).

10. Duforet-Frebourg, N., Luu, K., Laval, G., Bazin, E. & Blum, M. G. B. Detecting genomic signatures of natural selection with principal component analysis: Application to the 1000 Genomes data. Mol. Biol. Evol. 33, 1082–1093 (2016).

11. 1000 Genomes Project Consortium et al. A map of human genome variation from population-scale sequencing. Nature 467, 1061–1073 (2010).

12. 1000 Genomes Project Consortium et al. An integrated map of genetic variation from 1,092 human genomes. Nature 491, 56–65 (2012).

13. 1000 Genomes Project Consortium et al. A global reference for human genetic variation. Nature 526, 68–74 (2015).

14. Fairley, S., Lowy-Gallego, E., Perry, E. & Flicek, P. The International Genome Sample Resource (IGSR) collection of open human genomic variation resources. Nucleic Acids Res. 48, D941–D947 (2020).

15. Wiley, L. K. et al. Building a vertically integrated genomic learning health system: The biobank at the Colorado Center for Personalized Medicine. Am. J. Hum. Genet. 111, 11–23 (2024).

16. Suarez-Pajes, E., Díaz-de Usera, A., Marcelino-Rodríguez, I., Guillen-Guio, B. & Flores, C. Genetic Ancestry Inference and Its Application for the Genetic Mapping of Human Diseases. Int. J. Mol. Sci. 22, (2021).

17. Henn, B. M. et al. Cryptic distant relatives are common in both isolated and cosmopolitan genetic samples. PLoS One 7, e34267 (2012).

18. Hill, W. G. & Weir, B. S. Variation in actual relationship as a consequence of Mendelian sampling and linkage. Genet. Res. 93, 47–64 (2011).

19. Micheletti, S. J. et al. Genetic Consequences of the Transatlantic Slave Trade in the Americas. Am. J. Hum. Genet. 107, 265–277 (2020).

20. Gouveia, M. H. et al. Origins, Admixture Dynamics, and Homogenization of the African Gene Pool in the Americas. Mol. Biol. Evol. 37, 1647–1656 (2020).

21. Privé, F. et al. Portability of 245 polygenic scores when derived from the UK Biobank and applied to 9 ancestry groups from the same cohort. Am. J. Hum. Genet. 109, 373 (2022).

22. National Academies of Sciences, Engineering, and Medicine & Committee on the Use of Race, Ethnicity, and Ancestry as Population Descriptors in Genomics Research. Using Population Descriptors in Genetics and Genomics Research: A New Framework for an Evolving Field. (National Academies Press, 2023).

23. Stockwell, D. R. B. & Peterson, A. T. Effects of sample size on accuracy of species distribution models. Ecol. Modell. 148, 1–13 (2002).

24. Bean, W. T., Stafford, R. & Brashares, J. S. The effects of small sample size and sample bias on threshold selection and accuracy assessment of species distribution models. Ecography 35, 250–258 (2012).

25. Berahmand, K., Daneshfar, F., Salehi, E. S., Li, Y. & Xu, Y. Autoencoders and their applications in machine learning: a survey. Artificial Intelligence Review 57, 28 (2024).

26. Zhou, P., Xie, X., Lin, Z. & Yan, S. Towards Understanding Convergence and Generalization of AdamW. IEEE Trans. Pattern Anal. Mach. Intell. 46, 6486–6493 (2024).

27. Douze, M., et al. The Faiss library. *arXiv [cs.LG]* (2024).

28. Gaynor, S. M. et al. Yield of genetic association signals from genomes, exomes and imputation in the UK Biobank. Nat. Genet. (2024) doi:10.1038/s41588-024-01930-4.

29. Rieke, N. et al. The future of digital health with federated learning. NPJ Digit Med 3, 119 (2020).

30. Fernández, A., del Río, S., Chawla, N. V. & Herrera, F. An insight into imbalanced Big Data classification: outcomes and challenges. Complex & Intelligent Systems 3, 105–120 (2017).

31. Shi, Z. Improving k-Nearest Neighbors Algorithm for Imbalanced Data Classification. IOP Conf. Ser.: Mater. Sci. Eng. 719, 012072 (2020).

32. Arriaga-MacKenzie, I. S. et al. Summix: A method for detecting and adjusting for population structure in genetic summary data. Am. J. Hum. Genet. 108, 1270–1282 (2021).

